# Cell Size Controls Photosynthetic Capacity in a Mesoamerican and an Andean Genotype of *Phaseolus vulgaris* L

**DOI:** 10.1101/2024.02.13.580151

**Authors:** Andrew Ogolla Egesa, C. Eduardo Vallejos, Kevin Begcy

**Affiliations:** Environmental Horticulture Department, University of Florida, Gainesville, FL 32611, USA; Horticultural Sciences Department, University of Florida, Gainesville, FL 32611, USA; Plant Molecular and Cellular Biology Graduate Program, University of Florida, Gainesville, FL 32611, USA

**Keywords:** Carboxylation, common bean, gene pools, leaf anatomy, photosynthetic efficiency

## Abstract

The efficiency of CO_2_ flux in the leaf is hindered by a several structural and biochemical barriers which affect the overall net photosynthesis. However, the dearth of information about the genetic control of these features is limiting our ability for genetic manipulation. We performed a comparative analysis between a Mesoamerican and an Andean cultivar of *Phaseolus vulgaris* at variable light and CO_2_ levels. The Mesoamerican bean had higher photosynthetic rate, maximum rate of rubisco carboxylase activity and maximum rate of photosynthetic electron transport at light saturation conditions than its Andean counterpart. Leaf anatomy comparison between genotypes showed that the Mesoamerican bean had smaller cell sizes than the Andean bean. Smaller epidermal cells in the Mesoamerican bean resulted in higher stomata density and consequently higher stomatal conductance for water vapor and CO_2_ than in the Andean bean. Likewise, smaller palisade and spongy mesophyll cells in the Mesoamerican than in the Andean bean increased the cell surface area per unit of volume and consequently increased mesophyll conductance. Finally, smaller cells in the Mesoamerican also increased chlorophyll and protein concentration per unit of leaf area. In summary, we show that differential cell size controls the overall net photosynthesis and could be used as a target for genetic manipulation to improve photosynthesis.

**Highlight:** PhotosyntheUc performance comparison between a Mesoamerican and an Andean bean genotype showed higher rate at increased light and CO_2_ levels. Differences could be explained by variaUon in cell size.

## Introduction

Enhancing photosynthetic efficiency can improve plant performance and productivity (Baker *et al*., 2007; Cardona *et al*., 2018; Lin *et al*., 2022; Keller *et al*., 2024). We have a limited understanding of how genetic variation in anatomical, biochemical, and physiological architectures of the photosynthetic gas exchange apparatus impacts net photosynthesis (A_net_) (Baker et al. 2007; Sakoda et al. 2022). Nevertheless, recent studies have identified inter- and intra-specific phenotypic variation in photosynthetic gas exchange structures associated with adaptation to different environments (Tanaka 2019; Müller & Munné-Bosch 2021; Cackett et al. 2022; Sakoda et al. 2022). For instance, adaptation to a wide range of hydrological environments by species of *Banksia* are related to changes in morphological and anatomical characteristics that impact net assimilation (see review by Drake et al. 2013; and Harrison et al. 2020). Similarly, analysis of wild beans originated in central and south America indicated variabilities in photosynthetic nitrogen use efficiency, stomatal density, leaf chlorophyll content and protein solubility due to adaptation to local temperature and precipitation conditions (Gepts and Bliss, 1988; Koenig *et al*., 1990; Jonathan Lynch *et al*., 1992; Kami *et al*., 1995; Seher *et al*., 2014). Some of these modifications involved leaf size and photoperiod responses which could be associated with photosynthetic capacity (Schmutz *et al*., 2014). Thus, genetic variants could be exploited to improve the photosynthetic efficiency of different crops, particularly in the context of climate change.

The common bean (*Phaseolus vulgaris, L.)* is the most cultivated legume used for direct human consumption (Gepts, 2001), and it represents a significant component of the protein and carbohydrate caloric intake for over half a billion people worldwide (Siddiq et al. 2011; OECD 2019; Uebersax et al. 2022). Therefore, improving the productivity of common bean will have a significant global impact on food security. The potential for improvement is based on the extent of variation in this species. DNA sequence analysis revealed that Mesoamerica is the primary center of diversity of *P. vulgaris*, from which it radiated to the Andean region (Gaut, 2014). Furthermore, allele frequency analysis also indicated that beans were domesticated independently in each gene pool (Schmutz *et al*., 2014).

Several groups have matched the extent of genotypic diversity between the gene pools and wild and cultivated beans with the phenotypic diversity of traits associated with the photosynthetic activity. For instance, Lynch et al. (1992) reported extensive variation in leaf morphology, anatomy, biochemistry, and assimilation rates among a relatively large set of wild accessions from both gene pools. More recently, associations of certain DNA variants of beans with ecological niches independent of geographical distributions have been reported (Rodriguez *et al*., 2016). In addition, significant phenotypic differences in photosynthesis parameters between Mesoamerican and Andean cultivars have also been found (Sexton et al. 1997). These finding pointed out that these differences were responsible for the contrasting relative growth rates between the two groups.

Genetic characterization of the extant variation in photosynthesis-associated traits between Mesoamerican and Andean beans could enable genetic manipulations of the photosynthetic apparatus. In this study, we used two common beans that originated from separate domestication events, Calima domesticated in the Andean region and Jamapa domesticated in Mesoamerica. Thus, both genotypes present a unique opportunity to study their photosynthetic capacity. Towards this end, we examined the relationships between carbon assimilation responses to light and CO_2_ and their structural components with their photosynthetic capacity.

## Materials and Methods

### Plant materials and growth conditions

We selected for comparative analysis a representative genotype from each of the two *Phaseolus vulgaris L*. gene pools. Jamapa is a small, black-seeded landrace from Mesoamerica with an indeterminate growth habit, and Calima is a mottled large-seeded Andean bean with a determinate growth pattern. In addition, these genotypes exhibit contrasting photoperiod sensitivity (Bhakta *et al*., 2017), and are the parents of a recombinant inbred family (Bhakta et al., 2015).

Seeds from both genotypes were germinated in a 72-well nursery tray custom black (Nursery Supplies, FL) and transplanted at ten days into one-gallon black molded nursery cans. Three-week-old plants from each genotype were used for all the experiments. The media used in the nursery and transplanting was PRO-MIX HP Mycorrhizae planting media (Premier Horticulture, Canada). After transplanting, 17 g of Osmocote (N:P: K 18:6:12) were added to each pot. The greenhouse temperatures were maintained at 26±3 °C/ 20±3 °C day/night respectively. Irrigation was provided daily by applying water to field capacity.

### Physiological measurements

The uppermost completely expanded mature trifoliate leaf of each genotype was used to measure CO_2_ assimilation (A), transpiration rate (E), stomatal conductance to water vapor (g_sw_), intracellular CO_2_ (C_i_), and total conductance to CO_2_ (g_tc_), using the LI-COR Li-6800 machine (Begcy *et al*., 2019; LI-COR, 2023). The temperature inside the leaf chamber was maintained at 25 °C.

### CO_2_ conductance measurements

We exposed the mature trifoliate leaf of each genotype to three levels (200, 400, and 600 µmol mol⁻¹) of CO_2_ in the Li6800 leaf chamber using 1000 μmols m^2^ s^-1^ (PPFD) and 25 °C as set temperature. We then compared the patterns of leaf conductance using the measured values of A, E, g_sw_, C_i_, and g_tc_. We then used A, C_a_, and C_i_ values to estimate the CO_2_ conductance from outside into the leaves using the (Boyer and Kawamitsu, 2011) formula g_c_=(A/(C_a_-C_i_)).

### Light Response Curves (LRCs)

Responses to light were measured at 25°C and relative humidity of 60% ± 2% at two levels of CO_2_: ambient (400 µmol mol⁻¹) and elevated (600 µmol mol⁻¹) CO_2_. The light levels were gradually increased from 0 to 2000 μmols m^-2^ s^-1^ (PPFD) (Begcy *et al*., 2019).

### CO_2_ Response Curves (A−Ci)

CO_2_ response (A−Ci) curves were obtained at moderate PPFD (1000 μmol m^−2^s^−1^) for both bean genotypes using 12 replicates. The ambient CO_2_ (C_a_) was adjusted between 50 and 1200 µmol mol⁻¹. V_cmax_ and J_max_ were estimated using a modified Farquhar-von Caemmerer-Berry model as described in the plantecophys package (Duursma, 2015).

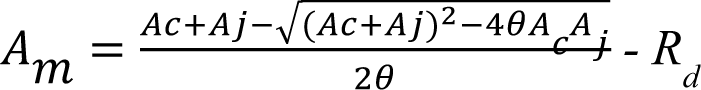

Where:

A_m_ = the hyperbolic minimum of A_c_ and A_j_ A_n_ = min (A_c_, A_j_) - R_d_

A_n_ = Net CO_2_ assimilation.

A_c_ = Photosynthesis rate when Rubisco activity is limiting.

A_j_ = Photosynthesis rate when RuBp –regeneration is limiting. R_d_ = the rate of mitochondrial respiration.

A_c_, the rubisco-limited photosynthesis rate was estimated as previously described (Duursma, 2015), and given by:

A_c_ = V_cmax_ (C_i_-Γ*) / [C_i_ + K_c_ (1 + O_i_/K_o_)]

Where V_cmax_ is the maximum rate of Rubisco activity, C_i_ and O_i_ are the intercellular concentrations of CO_2_ and O_2_, K_c_ and K_o_ are the Michaelis–Menten coefficients of Rubisco activity for CO_2_ and O_2_, respectively, and Γ* is the CO_2_ compensation point in the absence of mitochondrial respiration.

A_j_, the photosynthesis rate when ribulose-1,5-bisphosphate (RuBP)-regeneration is limiting was estimated as previously described (Laĭsk *et al*., 2009; Duursma, 2015), and according to:

A_j_ = (J/4) x (Ci - Γ*) /Ci + 2 Γ*)

Where J is the rate of electron transport which is related to incident photosynthetically active photon flux density, Q, by:

qJ^2^ -(aQ + J_max_) J + aQJ_max_ = 0 (when J < J_max_) where;

q = Is the quantum energy state.

a = absorptance by leaf photosynthetic pigments.

### Chlorophyll quantification

A set of leaf discs measuring 2.01 cm^2^ from fresh leaf tissue were harvested from each genotype. After fresh weight determination, discs were finely ground in liquid nitrogen and dissolved in four volumes of 100% of ice-cold acetone. The homogenate was brought up to 1 mL with 80% ice-cold acetone to and mixed by vortexing for 20 seconds. The mixture was centrifuged at 2000 g for 5 minutes, and the supernatant was obtained. Afterward, 150 μL of the chlorophyll extract was used to read the absorbance using a plate reader at 645nm and 663nm wavelengths to estimate chlorophyll a (ChlA) and chlorophyll b (ChlB), respectively. Total chlorophyll was calculated by the sum of Chlorophyll a and Chlorophyll b as described previously (Warren, 2008; Begcy *et al*., 2012).

### Total protein quantification

An additional set of leaf discs measuring 2.01 cm^2^ each from fresh leaf tissue were harvested for the total protein quantification. Discs were finely ground in liquid nitrogen and dissolved with equal volume (v/v) of the 2X Protein Extraction buffer (PE buffer: 0.1 M tris-HCl, pH 8; 2% SDS; 0.05 mL 1M DTT). Followed by the addition of 1X PE buffer to obtain a 1 mL sample-PE buffer mixture before further mixing by vortexing and progressing with the protein extraction. The mixture was then heated in a water bath at 100°C for 10 minutes, then allowed to cool at room temperature for 10 minutes, then pelleted at 20,000 g for 10 minutes at 23±1°C. 200 μL of the supernatant were transferred to new centrifuge tubes and mixed with 800 μL of 100% acetone. The mixture was centrifuged at 20,000 g at 23±1°C for 10 minutes and the supernatant was discarded. The pellet was allowed to dry at room temperature for 2 mins, then dissolved in 50 mL of 0.2 N NaOH and neutralized with an equal volume of 0.2 N HCl. The total protein content was determined using the colorimetric Bio-Rad Protein Assay Kit II (Bio-Rad Laboratories, CA). Seven dilutions of a protein standard containing 0 to 30 µg/mL of the total protein content were used. A standard curve was prepared each time the assay was performed. The absorbance at 595 nm (A595nm) was then measured with a microplate spectrophotometer (Epoch Microplate Spectrophotometer; BioTek, Winooski, VT), and the normalized absorbance values were plotted versus the mass concentration (µg of protein/mg of leaf tissue) as previously described (Kalaman *et al*., 2022).

### Stomatal density

To quantify stomatal density, leaf samples were prepared using the modified leaf peel method (Lawrence et al., 2018). In brief, intact leaves were carefully covered with clear adhesive tape on the abaxial and adaxial sides to obtain a leaf tissue peel with intact cuticle, epidermal, and guard cells. Before imaging, the peels were kept moist in 1% PBS and a small area was excised and mounted on a glass slide. Imaging was performed on a Leica compound microscope (Wetzlar, Germany) at a magnification of 10X. The microscope was fitted with Leica microsystems CMS camera calibrated with Leica Application Suite X LAS X (3.7.4.23463) for imaging. We used 25 μL of 5% propidium iodide to enhance boundaries of the epidermal and guard cells. The image J software (Abràmoff et al., 2004; Schneider et al., 2012) was used to determine stomatal density, and leaf epidermal cell sizes.

### Stomata and guard cell size estimation

To quantify stomata size, guard cells size, and fully open stomata aperture area, detached fresh leaf samples were incubated in 150 mL of stomata opening buffer (50 mM KCl, 10 mM MES-KOH, pH 6.2; (Lawrence *et al*., 2018), for a period of 30 mins. Following incubation, a section was excised and mounted on a glass slide. Imaging was performed on a Leica compound microscope (Wetzlar, Germany) at magnifications of 40X objective lens. The images obtained from each genotype were used to determine the size of stomata, guard cells, and full stomata aperture area using the image J software (Abràmoff *et al*., 2004; Schneider *et al*., 2012).

### Cell layer measurements

Fresh leaf samples from three-week-old plants from each genotype were used to prepare cross-sections by hand. Sections were mounted on glass slides using thin forceps. The sections were stained with 25 μL of 5% propidium iodide and imaged at 10X objective lens on a Leica compound microscope (Wetzlar, Germany). Images were used to count the number of cell layers using the Leica microsystems CMS camera calibrated with Leica Application Suite X LAS X (3.7.4.23463).

### Palisade and Mesophyll cell size characteristics

Fresh leaf samples from three-week-old plants were obtained from each genotype and used for cell isolation. The palisade and mesophyll cells were isolated using a modified leaf cell isolation protocol (Endo *et al*., 2016). Excised leaf discs (0.5-inch diameter) with their epidermal cell layer peeled off were incubated in 1.7 mL Eppendorf tubes containing 1 mL of ice-cold cell isolation enzyme buffer (75% (wt/vol) cellulase ‘Onozuka’ R-10, 0.25% (wt/vol) macerozyme R-10, 0.4 M, mannitol, 8 mM CaCl_2_ and 5 mM MES-KOH) for 20 minutes on a rotating Biometra OV4 Compact Line Hybridization Oven Incubator set at 24°C. Macerated discs were removed from the Eppendorf tubes before centrifugation at 200 g for 5 min at 4°C. The supernatant was discarded, then the isolated cells were gently re-suspended in 500 μL of the ice-cold wash buffer (2 mM MES,125 mM CaCl_2_,154 mM NaCl, 5 mM KCl) before another round of centrifugation at 200 g for 5 min at 4°C. The supernatant was discarded, and the pellets were resuspended in 100 μL of the cell flotation buffer (4 mM MES, 0.4 M mannitol, and 15 mM MgCl_2_ at pH 5.7) (Yoo *et al*., 2007; Nanjareddy *et al*., 2016), before imaging on a compound microscope at 40X objective lens.

Dimensions of the Isolated cells were obtained using the Image J software (Abràmoff *et al*., 2004; Schneider *et al*., 2012). Side projections of palisade cells were used to obtain the diameter (D), radius (r=D/2), and length (L). The volume of palisade cells was estimated as v = μr^2^L, and the surface area was estimated as SA = 2μr^2^ + 2μrL. Spongy mesophyll cell size was calculated by estimating an average radius of a sphere from three diameter estimates (d1, d2, d3) then the volume was estimated as V= 4/3μr^3^ and the surface area was estimated as SA = 4μr^2^.

### Statistical analysis

Differences between photosynthetic parameters, anatomical and physiological traits of the two genotypes were statistically tested using the Welch’s t-test (α = 0.05). The CO_2_ response data was subjected to Farquhar—von Caemmerer—Berry; the FvCB model for C_3_ photosynthesis as implemented by Duursma (2015). The parameter estimated (V_cmax_, J_max_, R_d_, CO_2_ levels for transitions from carboxylation limited to RuBP regeneration limited photosynthesis).

## Results

### Genotypes from the Andean and Mesoamerican gene pools display different photosynthetic performance under high light and CO_2_ conditions

To characterize the influence of adaptation on photosynthetic capacity in *P. vulgaris*, we used two common bean genotypes from the Andean (Calima) and Mesoamerican (Jamapa) gene pools. First, we subjected both genotypes to low and moderate light intensity, 600 and 1000 µmol m^-2^ s^-1^ Photosynthetic Photon Flux Density (PPFD), respectively (Fig. 1A-B). We did not find significant differences between Jamapa and Calima in their photosynthetic levels (A). However, the transpiration rate (E) and stomatal conductance (gsw) levels were significantly higher in Jamapa at moderate light levels (Fig. 1B). At a higher light intensity, specifically 1800 µmol m^-2^ s^-1^ (PPFD) (Fig. 1C) and 2000 µmol m^-2^ s^-1^ (PPFD) (Fig. 1D), Jamapa showed consistently statistically significant higher A, E and gsw than Calima. Based on their origin, these results suggest that both genotypes have been adapted to different light intensities, with Calima adapting to the low light while Jamapa to high light intensity.

**Figure 1.**
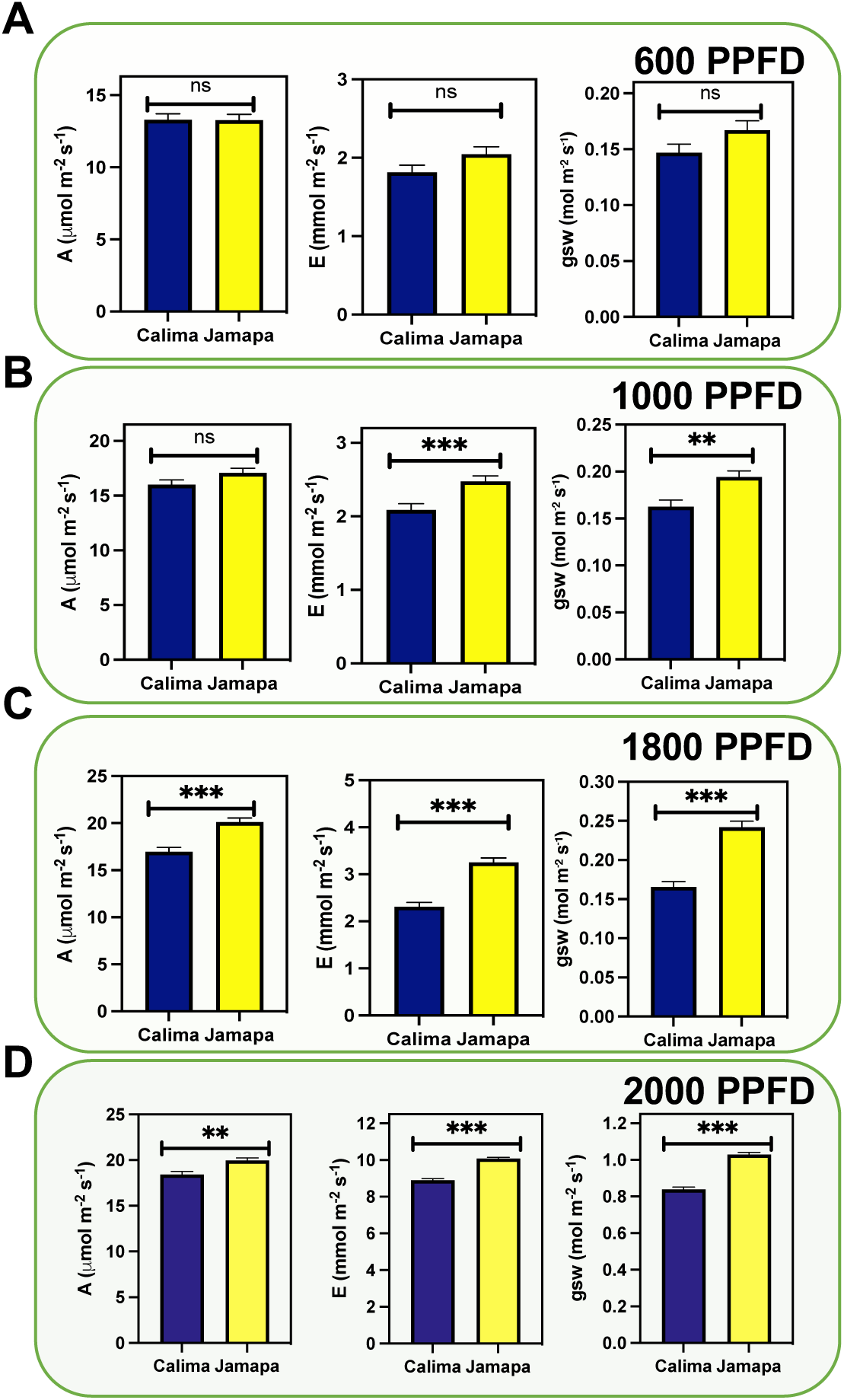
Photosynthetic capacity increases in the Mesoamerican genotype under higher light intensity. Photosynthetic gas exchange parameters (Assimilation [A], Transpiration [E], and Stomata conductance to water vapor [gsw]) were measured under different levels of light intensity at ambient CO2 (400 µmol m-2 s-1) represented by Photosynthetic Photon Flux Density (PPFD) levels. (A) 600 µmol m^-2^ s^-1^, (B) 1000 µmol m^-2^ s^-1^, (C) 1800 µmol m^-2^ s^-1^ and (D) 2000 µmol m^-2^ s^-1^ . n=20. Significant differences were calculated based on the Welch’s t-test at an alpha of 0.05. non-significant (ns) P > 0.05; **P ≤ 0.01, and *** P ≤ 0.001.

### Diurnal patterns of net assimilation in the Andean and Mesoamerican genotypes

To further characterize the impact on light adaptation of these two common bean genotypes, we analyzed their light compensation point and maximum rate of light-unlimited photosynthesis by using light curve measurements at ambient (400 µmol m^-2^ s^-1^) and elevated (600 µmol m^-2^ s^-1^) CO_2_ levels. We subjected both genotypes to increasing light intensity levels from 0 to 1800 µmol m^-2^ s^-1^ (Fig. 2). We used a modified hyperbolic function from light response curve (LRCs) using data from Calima and Jamapa to estimate the light compensation points (l*c*; x-axis intercept), the maximum photosynthetic rate at light-saturating conditions (V*lmax*; horizontal asymptote), the l*50* or PPFDs needed to attain 0.5 Vlmax and the quantum use efficiency (QUE= ΔA/ΔPPFD). In general, Jamapa exhibited a higher V*lmax* compared to Calima at both ambient and elevated CO_2_ (Fig. 2). The light compensation points (l*c*) for both genotypes were consistently lower in Calima (Fig. 2A-C). However, a shift from ambient to elevated CO_2_ resulted in a considerable drop in the light compensation point for both Calima and Jamapa (Fig. 2D-F).

**Figure 2.**
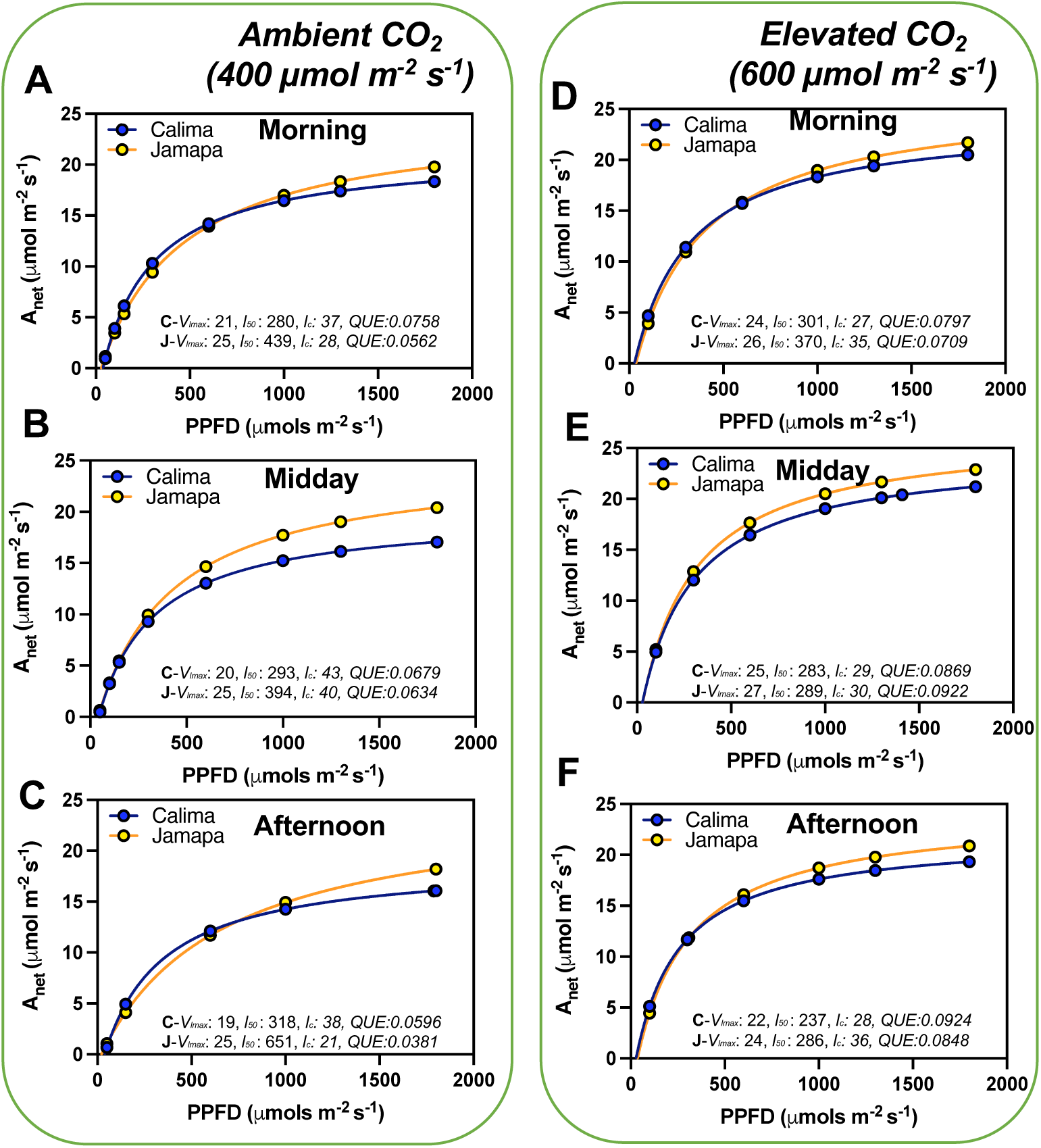
Diurnal photosynthetic light use efficiency characteristics are variable between Andean and Mesoamerican common bean genotypes. We used a modified hyperbolic function from light response curve (LRCs) using data to estimate the light compensation points (*l_c_*; x-axis intercept), maximum photosynthetic rate at light-saturating conditions (*V_lmax_*; horizontal asymptote), the *l_50_* or PPFDs needed to attain 0.5 *V_lmax_* and the quantum use efficiency (QUE= ΔA/ΔPPFD). These data were collected between 8 to 10 am (morning), 11 to 1pm (midday) and 2 to 4 pm (afternoon) (A-C) ambient CO_2_ (400 µmol m^-2^ s^-1^) and at (D-F) elevated CO_2_ (600 µmol m^-2^ s^-1^). n = 36.

### Higher carboxylation and electron transfer efficiencies in the Mesoamerican genotype

To further characterize CO_2_ conductance and carboxylation efficiencies of Calima and Jamapa, we subjected both genotypes to gradually changing levels of CO_2_ (A-Ci curve), using CO_2_ levels ranging from 50 to 1200 μmoles m^-2^ s^-1^. We estimated the maximum carboxylation (V*_cmax_*), Maximum rate of photosynthetic electron transport (J*_max_*), rate of dark respiration (R*_d_*), the amount of CO_2_ for the transition from ribulose-1,5-bisphosphate saturated to limited (RuBPsa-li) photosynthesis by fitting the A-Ci curves from photosynthesis data collected from Calima and Jamapa, where CO_2_ is the substrate in the reaction adopting the Farquhar—von Caemmerer—Berry; FvCB model for C3 photosynthesis as previously described (Duursma, 2015). Jamapa exhibited a higher Carboxylation efficiency (V*_cmax_*) and a higher electron transfer efficiency (J*_max_*) compared to Calima (Fig. 3A-F). These results were likely due to the prevailing efficient e-transfer rate or better CO_2_ conductance in Jamapa compared to Calima.

**Figure 3.**
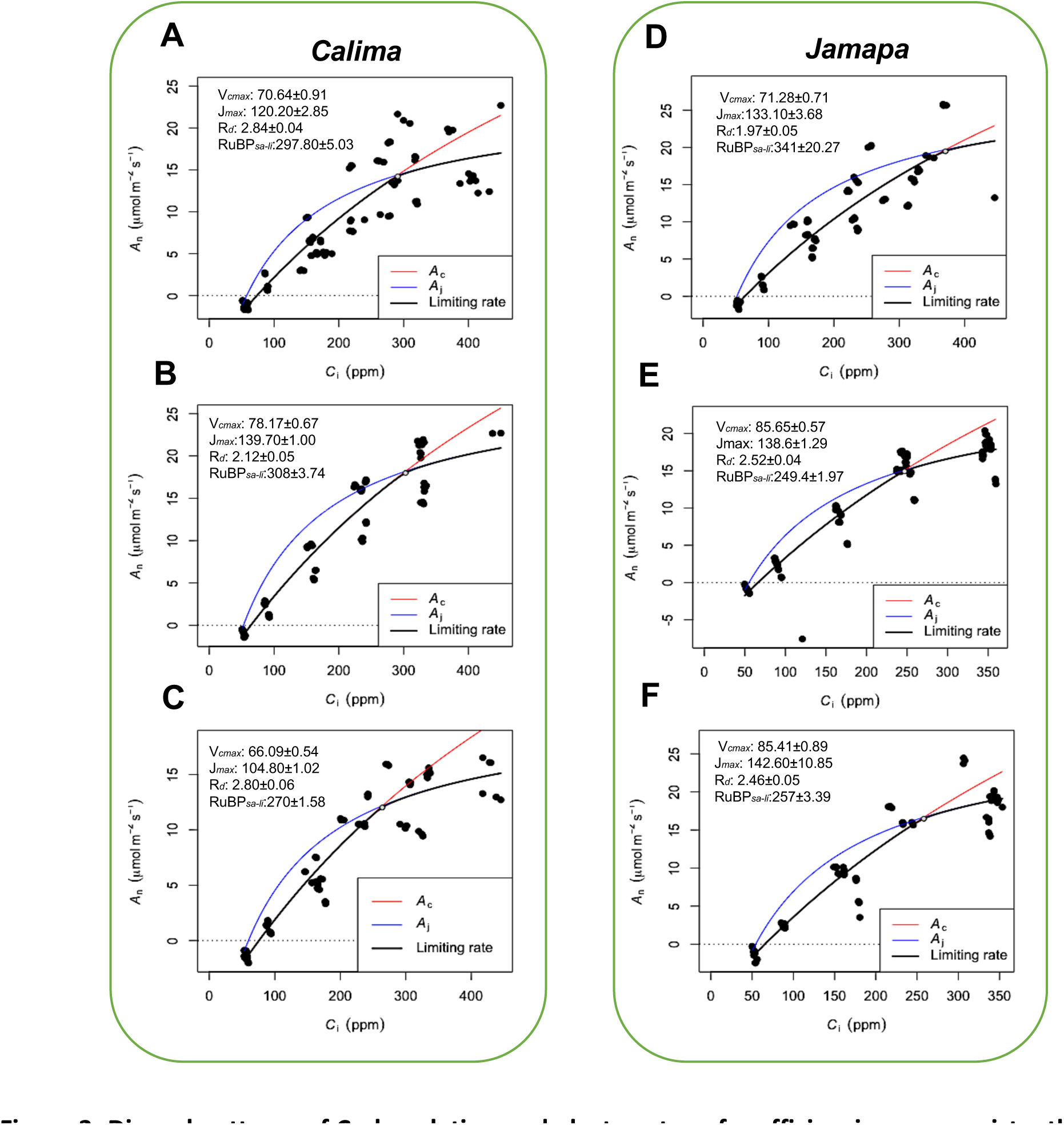
Diurnal patterns of Carboxylation and electron transfer efficiencies are consistently higher in the Mesoamerican common bean genotype and peak at different periods of the day. Maximum carboxylation (V*_cmax_*), Maximum rate of photosynthetic electron transport (J*_max_*), rate of dark respiration (R*_d_*), the amount of Co_2_ for the transition from ribulose-1,5-bisphosphate saturated to limited (RuBPsa-li) photosynthesis was estimated by fitting the A-Ci curves from photosynthesis data collected from Calima and Jamapa, where CO_2_ is the substrate in the reaction adopting the Farquhar—von Caemmerer—Berry. The FvCB model for C3 photosynthesis as described in Plantecophys package (Duursma, 2015). Estimated carboxylation electron transfer efficiencies, dark respiration, and Co_2_ levels for RuBP*_sa-li_* transition of Calima in the morning (A), midday (B), and the afternoon (C), and Jamapa in the morning (D), midday (E), and afternoon (F). Photosynthesis data was collected at 1000 μmol m^-2^ s^-1^ PPFD, temperature of 25°C, and Relative humidity of 60%. n =180.

Calima’s V*_cmax_* increased from the morning to midday and then dropped in the afternoon, and the J*_max_* values changed in a similar fashion. Dark respiration in Calima fluctuated between 2.12 and 2.84 µmol m^-2^ s^-1^ throughout the day with an apparent dip at mid-day (Fig. 3A-B). The internal CO_2_ concentration at which Calima switched from carboxylation-limited to RuBP-limited photosynthesis was stable in the morning and midday and decreased in the afternoon by 38 μmol mol^-1^ (Fig. 3). In contrast, Jamapa’s V*_cmax_* increased from morning to midday from 71.28 to 85.65 μmol m^-2^ s^-1^, and remained stable in the afternoon, while its J*_max_* had a net increase throughout the day (Fig. 3D-E). Jamapa’s dark respiration fluctuated between 1.97 in the morning to 2.52 μmol m^-2^ s^-1^ in the afternoon (Fig. 3D,F). Interestingly, Jamapa transitioned from RuBP-saturated to RuBP-limited CO_2_ assimilation at a C_i_ of 341 μmol mol^-1^ in the morning and a C_i_ of 249 μmol mol^-1^ at midday, then remained stable through the afternoon (Fig. 3D-F). In general, the V*_cmax_* and J*_max_* values of Jamapa were larger than those of Calima throughout the day, except for J*_max_* at midday (Fig. 3). However, the greatest difference between these genotypes was the daily dynamics of the transitions from RuBP saturated to RuBP limited photosynthesis at midday (Fig. 3B,E). The ratio of J_max_/V_cmax_ in Calima changed very little over the day, which was reflected in the narrow range of the C_i_’s at which the transition occurred. In Jamapa this ratio dropped from 1.9 to 1.6 throughout the day. Increased photosynthesis rate with increasing light in Jamapa compared to Calima suggested a better capture of light energy by the leaf which may have increased the availability of e^-^ and H^+^ to drive the photosynthesis reactions. These results indicate that Jamapa has a greater capacity to regenerate RuBP in the morning and that this capacity decays during the day, in contrast to Calima, which, comparatively, does not display such a dramatic change.

### Leaf anatomy as a predictor of photosynthetic efficiency in Calima and Jamapa

We hypothesized that some of the differences in photosynthetic characteristics observed between the two common bean genotypes could be explained due to their anatomical differences. To test this hypothesis, we performed comparative anatomical analyses of the leaf epidermis and the mesophyll.

First, we calculated the stomatal density using closed and opened stomata in Calima (Fig. 4A) and Jamapa (Fig. 4B). The stomatal density on the abaxial side of Jamapa leaves was 225±4.12/mm^2^ (Fig. 4C) and 66±2.08/mm^2^ (Fig. 4D) on the adaxial side. In contrast, the corresponding densities for Calima were 141±2.44/mm^2^ and 44±2.14/mm^2^ (Fig. 4C-D). Both genotypes exhibited comparable abaxial to adaxial density ratios – 3.4 and 3.2 for Calima and Jamapa, respectively – and consequently similar intergenotypic ratios (Fig. 4E). As a proxy to guard cell sizes, we measured the projected surface areas. On the abaxial side, Calima’s guard cells (204±1.35 μm^2^) were 15% larger than those of Jamapa (176±0.98 μm^2^) (Fig. 4F). On the adaxial side of the leaf, the estimated surface area of Calima’s guard cells (211.0±7.42 μm^2^) was not significantly different from those of Jamapa (232±8.56 μm^2^) (Fig. 4G). Furthermore, the apertures of fully open Calima stomata (76±0.81 μm^2^) on the abaxial side were 37% larger than those of Jamapa (55±0.51μm^2^) (Fig. 4H). However, on the adaxial side the stomata apertures were not significantly different, Calima 44.6±2.99 μm^2^ and Jamapa 47.32±4.48 μm^2^, data that agree with the guard cell sizes (Fig. 4I). Moreover, the stomata size was significantly larger in Calima than Jamapa in the abaxial size (Fig. 4J), but not in the adaxial size (Fig. 4K).

**Figure 4.**
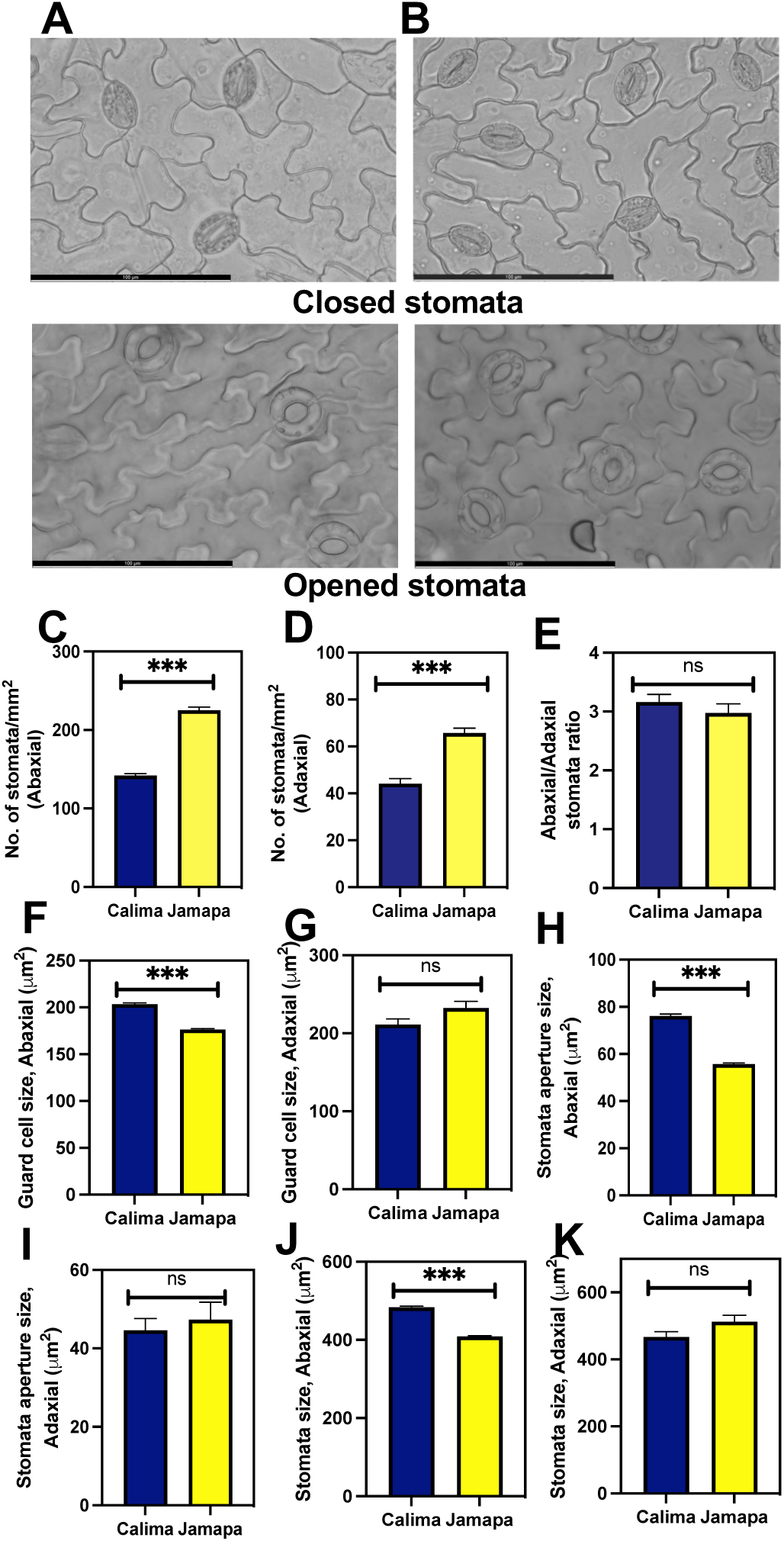
Andean common bean genotype exhibits lower stomatal density per unit area than its Mesoamerican counterpart. Light microscopy images of (A) Calima and (B) Jamapa depicting closed and opened stomata. Number of stomata on the (C) abaxial and (D) abaxial sides of the leaf. (E) Ratio of stomata per unit area on the abaxial to the adaxial side. Guard cell size on the (F) abaxial and the (G) adaxial side of the leaf. Open stomata aperture size on the (H) abaxial and the (I) adaxial side. Stomata size on the (J) abaxial and the (K) adaxial sides. Fresh mature leaves from three-week-old plants were incubated in 150 mL of stomata opening buffer for 30 mins to open the stomata. . Significant differences were calculated based on the Welch’s t-test at an alpha of 0.05. ns P > 0.05; * P ≤ 0.05, **P ≤ 0.01, and *** P ≤ 0.001. n = Abaxial: 346, Adaxial: 340. Scale bar = 100 μm.

To further elucidate whether the differences in photosynthetic capacity could be explained by their anatomical differences. We measured epidermal, palisade and mesophyll cells from both genotypes (Fig. 5A). First, we analyzed the size of the epidermal pavement cells (Fig. 5B). The average surface area of abaxial and adaxial pavement cells of Jamapa leaves was 1502±31 and 2940±82 μm^2^ and those of Calima 1674±32 and 3493±78 μm^2^, respectively (Fig. 5B-C). Thus, Calima pavement cells are 11 to 18% larger than Jamapa cells. Furthermore, we quantified the number of epidermal cells in the abaxial (Fig. 5D) and the adaxial size (Fig. 5E). Jamapa had higher quantity in both. An anatomical normalization or calculation of the number of pavement cells per stoma shows while Jamapa has 3 pavement cells per stoma on the abaxial side and 5 pavement cells per stoma on the adaxial (Fig. 5F-G), the corresponding ratios in Calima are 4 and 6 (Fig. 5F-G). In summary, physically and anatomically, Jamapa has a higher stomatal density than Calima.

**Figure 5.**
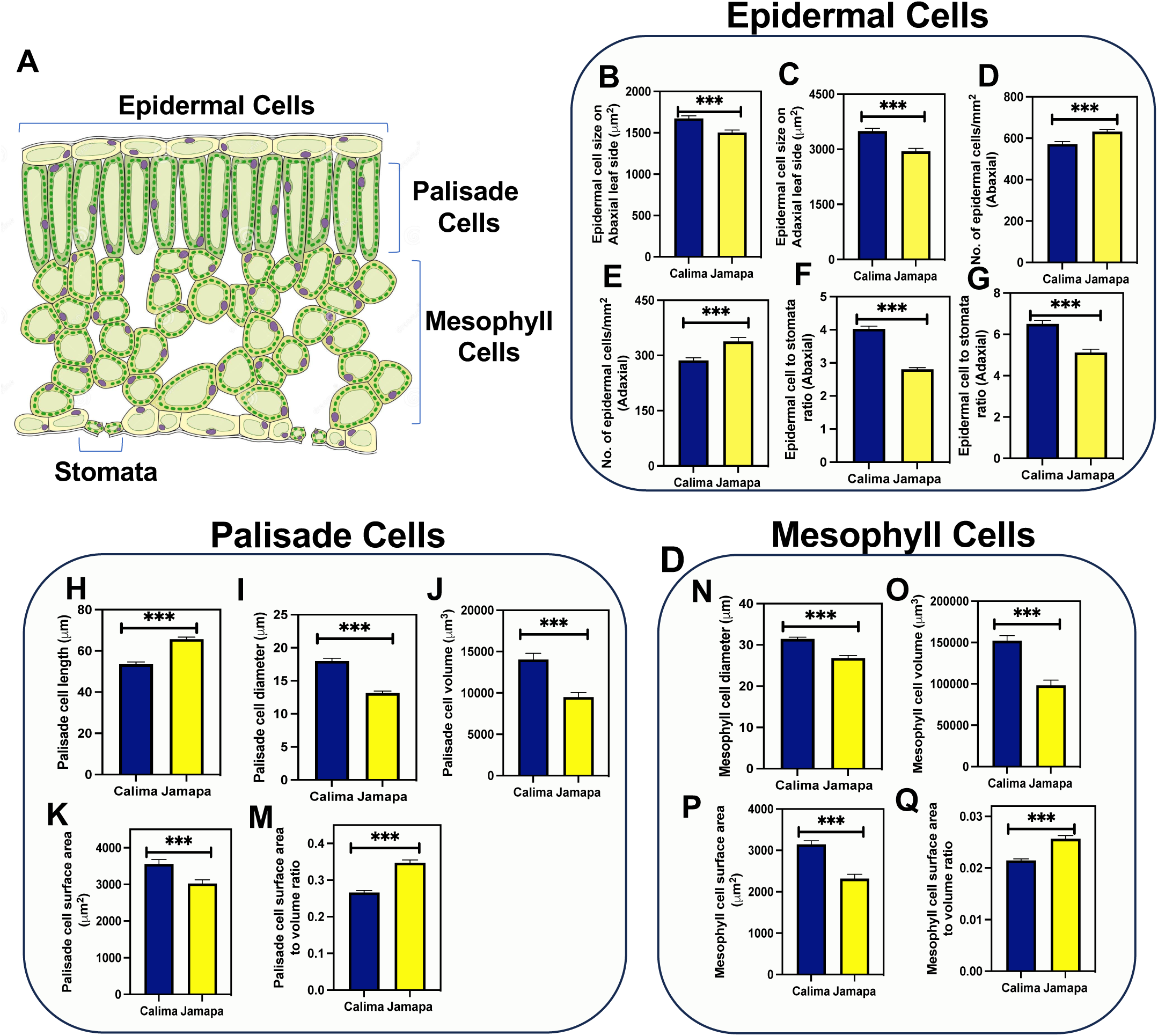
Epidermal, palisade, and mesophyll cell size is different between the Andean and the Mesoamerican genotypes. (A) Schematic representation of a cross section of a leaf indicating the location of stomata, epidermal, palisade and mesophyll cells. Epidermal cell sizes on the (B) abaxial and (C) abaxial sides. Number of epidermal cells on the (D) abaxial and (E) adaxial sides. Epidermal cells to stomata ratio on the (F) abaxial and (G) adaxial side in Calima and Jamapa. Palisade cells (H) length, (I) diameter and (J) volume. Palisade cells surface area (K) and surface area to volume ratio (M). Mesophyll cells (N) diameter, (O) volume and total surface area (P). Lateral projections of palisade cells were used to obtain the diameter (D), radius (r=D/2), and length (L). The volume of palisade cells was estimated as v = μr^2^L, and the surface area was estimated as SA = 2μr^2^ + 2μrL. Spongy mesophyll cell size was calculated by estimating the average radius of a sphere. The surface area was estimated as SA = 4μr^2^. Significant differences were calculated based on the Welch’s t-test at an alpha of 0.05. ns P > 0.05; * P ≤ 0.05, **P ≤ 0.01, and *** P ≤ 0.001. n = 45.

Examination of a cross-section of the leaf blade showed that the two genotypes had three layers of spongy parenchyma cells arranged below a single palisade cell layer (Supplementary Fig. S1). We isolated leaf palisade and spongy parenchyma cells and measured their lateral projections and used them to obtain first-order approximations of their cell volume and surface area. Calima palisade cells (53.52±1.10 µm) were shorter than Jamapa cells (65±0.96 µm) (Fig. 5H), but significantly wider (17±0.42 µm) than Jamapa cells (13.14±0.3 µm) (Fig. 5I). These dimensions were used to estimate cell volumes (Fig. 5J), and surface areas (Fig. 5K) assuming palisade cells as cylindrical and spongy parenchyma cells as spherical bodies. The average volume of Calima palisade cells (14038±760.6 µm^3^) was a 32% larger than those of Jamapa (9482±544.1µm^3^) (Fig. 5J), and the average surface area for Calima cells (3,556±122.4 µm^2^) was 14.99 % larger than that of Jamapa cells (3023±102.8 µm^2^) (Fig. 5K). Using these values, we calculated that Jamapa cells have 23 % more surface area than Calima cells per unit of volume in the palisade cells (Fig. 5M).

On the other hand, the average diameter of spongy parenchyma cells was larger in Calima (31.45±0.43 µm) than in Jamapa (26.79±0.63 µm) (Fig. 5N). Similarly, the average volume of spongy parenchyma cells of Calima (152037±6624 µm^3^) was 35% larger than those of Jamapa cells (98185±6421 µm^3^) (Fig. 5O), and the surface area of in Calima cells (3,146±85.50 µm^2^) was 26% greater than that of Jamapa cells (2,318±104.8 µm^2^) (Fig. 5P). Similarly, Jamapa cells have 19 % more in the spongy parenchyma than Calima cells per unit of volume (Fig. 5Q). Collectively, these results indicated that mesophyll cells in Jamapa have a larger surface area for CO_2_ diffusion than those of Calima.

### Effects of anatomical differences on physiological parameters

We measured the gas exchange parameters at three atmospheric CO_2_ concentrations (low C_a_ = 200, ambient C_a_ = 400, and high C_a_ = 600 μmol mol^-1^) to assess the effect of the anatomical differences in gas exchange characteristics (Table 1). Both genotypes displayed similar responses to the different C_a_ levels. Stomatal conductance to water vapor (g_sw_) increased when C_a_ was raised from low to ambient CO_2_, but decreased when C_a_ reached the highest level. Changes in E mirrored changes in stomatal conductance to water vapor (g_sw_). However, Jamapa displayed statistically significant greater g_sw_ and higher E rates than Calima at all C_a_ levels.

**Table 1.** Leaf assimilation and conductance in Calima and Jamapa. Conductance to CO_2_ (g_c_) was calculated using the Boyer and Kawamitsu (2011) formula g_c_=(A/Ca-Ci). Mesophyll conductance was calculated as the difference between the total conductance to CO_2_ (gtc) and the estimated CO_2_ conductance (g_c_). n = 60. For each parameter, a different letter within a column indicates significant differences based on the Welch’s t-test at an alpha of 0.05 comparing both genotypes. Same letter indicates no significant differences.

| Parameter | Genotype | 200 $\mu\text{mol mol}^{-1} \text{CO}_2$ | 400 $\mu\text{mol mol}^{-1} \text{CO}_2$ | 600 $\mu\text{mol mol}^{-1} \text{CO}_2$ |
| --- | --- | --- | --- | --- |
| A ($\mu\text{mol m}^{-2} \text{s}^{-1}$) | Calima | 6.20±0.23<sup>b</sup> | 15.42±0.48<sup>b</sup> | 17.81±0.45<sup>a</sup> |
| | Jamapa | 8.04±0.27<sup>a</sup> | 17.16±0.32<sup>a</sup> | 18.22±0.37<sup>a</sup> |
| C<sub>i</sub> ($\mu\text{mol mol}^{-1}$) | Calima | 161.10±1.14<sup>b</sup> | 306.48±3.67<sup>b</sup> | 458.72±5.28<sup>b</sup> |
| | Jamapa | 165.01±1.27 | 327.44±3.75 | 494.80±6.47<sup>a</sup> |
| $g_{sw}$ ($\mu\text{mol m}^{-2} \text{s}^{-1}$) | Calima | 0.34±0.01<sup>b</sup> | 0.38±0.02<sup>b</sup> | 0.28±0.02<sup>b</sup> |
| | Jamapa | 0.51±0.02<sup>a</sup> | 0.65±0.03<sup>a</sup> | 0.50±0.03<sup>a</sup> |
| E ($\text{mmol m}^{-2} \text{s}^{-1}$) | Calima | 4.14±0.15<sup>b</sup> | 4.45±0.2<sup>b</sup> | 3.47±0.18<sup>b</sup> |
| | Jamapa | 5.64±0.17<sup>a</sup> | 6.51±0.25<sup>a</sup> | 5.33±0.28<sup>a</sup> |
| $g_{tc}$ ($\text{mol m}^{-2} \text{s}^{-1}$) | Calima | 0.20±0.0077<sup>b</sup> | 0.22±0.0126<sup>b</sup> | 0.17±0.0098<sup>b</sup> |
| | Jamapa | 0.29±0.0097<sup>a</sup> | 0.36±0.017<sup>a</sup> | 0.28±0.0178<sup>a</sup> |
| $g_c$ ($\text{mol m}^{-2} \text{s}^{-1}$) | Calima | 0.18±0.0064<sup>b</sup> | 0.20±0.0107<sup>b</sup> | 0.15±0.0083<sup>b</sup> |
| | Jamapa | 0.26±0.0088<sup>a</sup> | 0.31±0.01<sup>a</sup> | 0.24±0.013<sup>a</sup> |
| $g_m$ ($\text{mol m}^{-2} \text{s}^{-1}$) | Calima | 0.02313±0.0016<sup>b</sup> | 0.02189±0.0016<sup>b</sup> | 0.0173±0.0016<sup>b</sup> |
| | Jamapa | 0.03752±0.0022<sup>a</sup> | 0.04785±0.0033<sup>a</sup> | 0.04669±0.0046<sup>a</sup> |
| WUE ($\mu\text{mol mmol}^{-1}$) | Calima | 1.50 | 3.46 | 5.13 |
|  | Jamapa | 2.63 | 2.63 | 3.41 |

Like g_sw_, stomatal conductance to CO_2_ (g_sc_) increased in both genotypes as C_a_ increased from low to mid-level, but dropped significantly when C_a_ reached the highest level (Table 1). Unlike E rates, net CO_2_ assimilation rates (A) increased to the highest level in both genotypes as CO_2_ levels increased (Table 1). Jamapa displayed statistically significantly higher A_n_ rates than Calima at low and mid C_a_ levels, however, these differences disappeared at the highest C_a_ as A_n_ reached saturation in both genotypes, although Jamapa showed statistically higher C_i_ values than Calima at all C_a_ levels. The intercellular CO_2_ concentrations (C_i_) increased in both genotypes proportionally to the increases in C_a_ maintaining a Ci/Ca ratio around 0.8. However, this ratio was lower in Calima than in Jamapa, particularly at the mid and high C_a_ levels. In summary, the response pattern of E, C_i_, A_n_, g_sw_, and g_cs_ to different levels of C_a_ was similar in both genotypes, but Jamapa displayed significantly higher values throughout. These differences can be explained largely by the fact that the total stomata aperture per unit of leaf area of Jamapa is 15 % larger than that of Calima. However, Jamapa’s g_sw_ and g_sc_ exceed those of Calima’s by up to 44 to 78%. The disproportionality between total stomatal aperture per unit of leaf area and the estimated conductance strongly suggests other functional differences in addition to those of their epidermal anatomies.

An analysis of leaf-level water use efficiency (WUE) under current CO_2_ level (400 μmol mol^-1^) and possible future higher level (600 μmol mol^-1^) showed that Calima currently outperforms Jamapa by about 30%, an advantage that could increase to 50% under the high CO_2_ level (Table 1).

We used the FvCB model to estimate mesophyll conductances (g_m_). Jamapa’s g_m_ was from 60 to 170 % larger than that of Calima at all C_a_ levels. Differences in g_m_ could be explained in part by the fact that cell surface area per unit of volume of Jamapa’s palisade and spongy parenchyma cells are 28 and 17 %, respectively, larger than those of Calima cells. Interestingly, Jamapa’s g_m_ increased by 30 % when C_a_ increased from 200 to 400 μmole/mole, while Calima’s dropped by 25% (Table 1).

### Jamapa has higher chlorophyll and total protein content per unit area but carboxylation reactions in Calima are more efficient

We measured chlorophyll and protein content per unit of leaf area to investigate whether the differences in cell size between the genotypes could give rise to differences in the density of components of the photosynthetic apparatuses; these differences, if any, could also explain to some extent differences in photosynthetic capacities. Protein analysis indicated that there were statistically significant differences between Calima (399.9 mg/m^2^) and Jamapa (630.10 mg/m^2^) (Table 2). We found that both genotypes had similar chlorophyll a/b ratios: Calima (2.82) and Jamapa (2.65). However, the total chlorophyll content in Jamapa (169.02 mg/m^2^) was significantly higher than Calima’s (149.41 mg/m^2^) (Table 2).

**Table 2.** Leaf chlorophyll and protein characteristics between Calima and Jamapa. Chlorophyll and total protein were measured in mature fully expanded leaves (n = 8). A different letter within a column indicates significant differences for each parameter based on Welch’s t-test at an alpha of 0.05 comparing both genotypes. Same letter indicates no significant differences.

| Genotype | chlorophyll a<br>(mg/m<sup>2</sup>) | chlorophyll b<br>(mg/m<sup>2</sup>) | chlorophyll a/b<br>ratio | Total chlorophyll<br>(mg/m<sup>2</sup>) | Total protein<br>(mg/m<sup>2</sup>) |
| --- | --- | --- | --- | --- | --- |
| Calima | 108.0± 3.76<sup>b</sup> | 41.42±2.35<sup>a</sup> | 2.82± 0.19<sup>a</sup> | 149.41±0.76<sup>b</sup> | 399.90±25.84<sup>b</sup> |
| Jamapa | 122.3± 3.95<sup>a</sup> | 46.74±1.86<sup>a</sup> | 2.65± 0.06<sup>a</sup> | 169.02±1.13<sup>a</sup> | 630.10±50.65<sup>a</sup> |

After estimation of the total leaf chlorophyll and the total protein content corresponding to the unit area used in A_net_ calculation, we fitted a modified hyperbolic function from the light response curve (LRCs) data from Calima and Jamapa to estimate, first, the amount of CO_2_ assimilated by chlorophyll unit (Fig. 6A) and the amount of CO_2_ assimilated per total protein (Fig. 6B) unit for given light intensity levels. Jamapa showed a higher V_lmax_ and l_50_ compared to Calima. The estimated values for V_lmax_ and l_50_ after normalizing them by total chlorophyll (V_lmax_:129 and l_50_:264 for Calima and V_lmax_: 146 and l_50_: 409 for Jamapa). Contrary, when we normalized the A_net_ by total protein (Fig. 6B), Calima had higher V_lmax_. Furthermore, the estimated l_c_ remained similar between chlorophyll and total proteins (Fig. 6), but Calima had a higher l_c_ (41) compared to Jamapa (35). Finally, the QUE in Calima was 0.49 after accounting for the total chlorophyll and 0.18 after accounting for the total protein (Fig. 6). In Jamapa, the QUE was 0.36 after normalizing it for the total chlorophyll and 0.10 after normalizing it for the total protein (Fig. 6). In comparison, the QUE values were higher in Calima compared to Jamapa.

**Figure 6.**
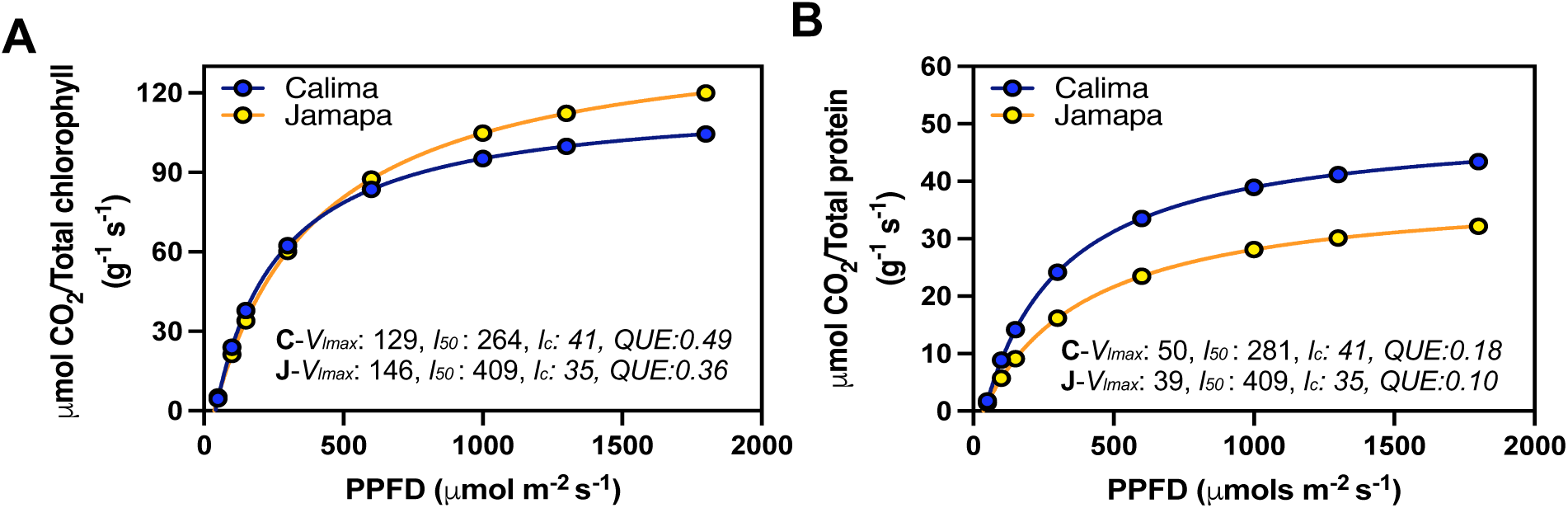
Photosynthetic light use efficiency after accounting for differences in chlorophyll and protein content between Andean and Mesoamerican common bean genotypes. LRCs from photosynthesis estimated per unit chlorophyll (A) and unit protein (B) from Calima and Jamapa. Fitting a modified hyperbolic function from light response curve (LRCs) data from Calima and Jamapa to estimate the light compensation points (*l_c_*; x-axis intercept), maximum photosynthetic rate at light-saturating conditions (*V_lmax_*; horizontal asymptote), the *l_50_* or PPFDs needed to attain 0.5 *V_lmax_ and an* estimated quantum use efficiencies (QUE) by calculating the first derivative of the light function at *l_c._* These estimates were from data collected between 8 am to 2.00 pm during the day at ambient CO_2_ 400 µmol m^-2^ s^-1^. n=36.

Considering this difference, we calculated the maximum rate of CO_2_ fixation based on protein content. Accordingly, Calima’s V_cmax_ rate (167.8 μmole g^-1^ sec^-1^) was 26% greater than Jamapa’s (125.0 μmole g^-1^ sec^-1^) (Table 3). However, when results were expressed on a leaf area basis, Jamapa exceeded Calima by 12%. Regarding the electron transport efficiency J_max_, Jamapa (251.9 μEq g^-1^ sec^-1^) retained significantly higher values than Calima (137.9 μEq g^-1^ sec^-1^). The rate of dark respiration remained higher in Calima (4.72 μmole g^-1^ s^-1^) compared to Jamapa (3.31 μmole g^-1^ s^-1^). At the same time, the internal CO_2_ concentration at which Calima switched from carboxylation-limited to RuBP-limited was lower in Calima (55.56 μmol mol^-1^) and significantly higher in Jamapa (259 μmol mol^-1^) (Table 3). These results indicated that the carboxylation reactions in Calima are more efficient than those in Jamapa, while photosynthetic electron transport efficiency is higher in Jamapa.

**Table 3.** Carboxylation and electron transfer efficiencies of an Andean and a Mesoamerican common bean genotypes on total protein per unit area basis. The carboxylation efficiency on a total protein basis was estimated at moderate light (1000 µmol m^-2^ s^-1^ PPFD). Maximum carboxylation (V_cmax_), Maximum rate of photosynthetic electron transport (J_max_), rate of dark respiration (R_d_) was estimated by fitting the A-Ci curves from photosynthesis data collected from Calima and Jamapa, where CO_2_ was the substrate in the reaction adopting the Farquhar—von Caemmerer—Berry; FvCB model for C3 photosynthesis as implemented by (Duursma, 2015). Photosynthesis data was obtained under a standard light intensity of 1000 μmol m^-2^ s^-1^ PPFD, temperature of 25°C, and Relative humidity of 60%. (n = 9). For each parameter, same letter indicates significant differences based on the Welch’s t-test at an alpha of 0.05 comparing both genotypes.

| Genotype | A-Ci |  |  |  |
| --- | --- | --- | --- | --- |
| | $V_{\text{cmax}}$<br>($\mu\text{mol g}^{-1} \text{s}^{-1}$) | $J_{\text{max}}$<br>($\mu\text{Eq g}^{-1} \text{sec}^{-1}$) | $R_d$<br>($\mu\text{mol g}^{-1} \text{s}^{-1}$) | $\text{RuBP}_{\text{sa-li}}$<br>($\mu\text{mol mol}^{-1}$) |
| Calima | 167.8±1.44<sup>a</sup> | 137.9±2.68<sup>a</sup> | 4.72±0.19<sup>a</sup> | 55.56±4.01<sup>b</sup> |
| Jamapa | 125.0±0.26<sup>b</sup> | 251.9±1.12<sup>b</sup> | 3.31±0.03<sup>b</sup> | 259.0±1.17<sup>a</sup> |

## Discussion

Our analysis showed that the Mesoamerican bean Jamapa has a V_lmax_ that is 22% greater than that of the Andean bean Calima. The V_cmax_ is 12% greater in Jamapa than in Calima. However, when the V_cmax_ is expressed on the basis of the total protein instead of leaf area, the Jamapa’s advantage is nullified, and Calima has more efficient carboxylation reactions. These comparisons strongly suggest that these genotypes have comparable photochemical capacities, but the structural differences that control the protein and chlorophyll content per unit of leaf area have a significant effect on their photosynthetic capacities.

Evidence from other studies has indicated the importance of the stomatal density in controlling the flow of CO_2_ into the intracellular spaces (Tomás *et al*., 2013; Harrison *et al*., 2020). Higher stomatal density promotes a better stomatal conductance (g_s_) (Harrison *et al*., 2020), mesophyll conductance to CO_2_ (Kaiser *et al*., 2019) and are essential in fine-tuning the CO_2_ flow and plant water balance. In addition, larger stomata apertures have been observed to lag in the opening and closing (Drake *et al*., 2013). Our data shows that differences in cell size between the genotypes appear to have the most consequential effect among other structural differences that affect photosynthetic capacity. Calima clearly has larger pavement cells than Jamapa which in effect lower the stomatal densities on the abaxial and adaxial sides of the leaf (Fig. 4). The larger guard cells and stoma opening of Calima are not able to counteract the low density caused by the larger pavement cells. In summary, total stomatal opening in Jamapa’s abaxial and adaxial leaf sides are 13 and 8% greater than in Calima’s, respectively (Fig. 4).

As expected, the differences in cell sizes and stomatal apertures had a significant effect on stomatal conductance to water vapor and CO_2_. As a result of the higher stomatal conductance, Jamapa displayed greater rates of transpiration and CO_2_ assimilation than Calima. However, by the same token, Calima displayed greater WUE than Jamapa. This phenomenon should not be overlooked in light of climate change upon us where higher CO_2_ levels and temperatures are expected, conditions in which Calima is likely to have an advantage.

Cell sizes are critical in mesophyll conductance in the diffusion the CO_2_ across several membranes into the chloroplast for CO_2_ fixation (Flexas *et al*., 2008; Tomás *et al*., 2013; Théroux-Rancourt and Gilbert, 2017; Elferjani *et al*., 2021; Momayyezi *et al*., 2022*a*).

We also detected significant cell size differences in parenchyma and spongy mesophyll cells. Overall, Jamapa’s smaller cells resulted in larger cellular surface area per unit of leaf volume than what was measured for Calima (Fig. 5). This meant that Jamapa mesophyll cells provide a larger area for CO_2_ diffusion into the cells than Calima cells, a phenomenon that is reflected in Jamapa’s larger mesophyll conductance. These results also suggest that the difference in chlorophyll content per unit of leaf area may be due to the differences in the size of mesophyll cells, provided the number and size of chloroplasts in the cells of each genotype are very similar.

There are differences in the size of pavement and guard cell between the abaxial and adaxial sides of the leaf in both genotypes. However, we notice there is a lack of proportionality between genotypes. This result suggests that the developmental controls of the two sides may be independent to some extent. In addition, the differences in the number of pavement cells per stomata between genotypes also suggest another developmental polymorphism between the genotypes. Larger and thicker leaves have been previously linked to an adaptation to lowlands in Juglans (Momayyezi *et al*., 2022*b*). Thus, longer palisade cells in Jamapa might be an adaptation to lower attitudes in the Mesoamerican region. Such thick leaves offer a double advantage of utilizing more light energy through increased chloroplasts and a higher mesophyll conductance (Momayyezi *et al*., 2022*b*). Therefore, leaf anatomical structure in Jamapa facilitates the efficient utilization of increasing light intensity, resulting in higher net photosynthesis. Therefore, while larger palisade and mesophyll cells surface area to volume ratio appear to be an adaptation to high light intensity, the opposite is suitable for low light intensity in common beans.

A comparative analysis of Mesoamerican and Andean cultivars detected significant differences in organ size (Sexton *et al*., 1997). A cluster analysis of 427 bean genotypes from both gene pools documented that the main difference between the pools is yield, with the Mesoamerican lines excelling over the Andean lines (Amongi *et al*., 2023). The results presented here strongly suggest that the yield differences between the gene pools are most likely due to differences in photosynthetic capacity as influenced by their differences in anatomic characters. Furthermore, the availability of a genotyped recombinant inbred family produced between Jamapa and Calima (Bhakta *et al*., 2015), will facilitate the genetic analysis of cell size and testing of the hypothesis that cell size exerts significant control over photosynthetic performance.

## Acknowledgments

We thank Simon Riley from the IFAS statistics consulting unit for guidance on the statistical analysis for the manuscript.

## Author Contributions

K.B. and C.E.V. Designed and supervised the experiments. A.O.E. Performed the experiments. A.O.E., C.E.V., and K.B. analyzed the data and wrote the manuscript. All authors read and approved the manuscript.

## Data Availability

All data collected for this study is included within the manuscript and its supporting information files.

## Abbreviations

A (μmol m^-2^ s^-1^): Assimilation rate
A_n_/A_net_ (μmol m^-2^ s^-1^): Net photosynthesis/Net assimilation C_i_ (μmol mol^-1^)
C_a_ (μmol mol^-1^): CO_2_ in the external environment E (μmol m^-2^ s^-1^)
g_sw_ (mol m^-2^ s^-1^): Stomatal conductance to water vapor g_tc_ (mol m^-2^ s^-1^)
g_s_ (mol m^-2^ s^-1^): Stomata conductance to CO_2_/g_c_ CO_2_ conductance into the leaf g_b_ (mol m^-2^ s^-1^)
g_m_ (mol m^-2^ s^-1^): Mesophyll conductance
V_lmax_ (μmol m^-2^ s^-1^): Maximum photosynthetic rate at light-saturating conditions l_50_ (μmol m^-2^ s^-1^)
V_cmax_ (μmol m^-2^ s^-1^): Maximum rate of Rubisco carboxylase activity
J_max_ (μEq m^-2^ sec^-1^): Maximum rate of photosynthetic electron transport R_d_ (μmol m^-2^ s^-1^)

